# Simultaneous *in vivo* imaging of Ca^2+^ signals in periarteriolar cholinergic axonal varicosities and arteriole diameter changes in the mouse cerebral cortex

**DOI:** 10.64898/2026.04.30.721808

**Authors:** Nobuhiro Watanabe, Harumi Hotta

**Affiliations:** Department of Autonomic Neuroscience, Tokyo Metropolitan Institute for Geriatrics and Gerontology

**Keywords:** basal forebrain cholinergic neuron, axonal varicosity, cerebral arteriole, somatosensory stimulation, two-photon imaging, *in vivo*

## Abstract

Basal forebrain cholinergic neurons project widely to the cerebral cortex and participate in cerebrovascular regulation. Although cholinergic axons are distributed around the cerebrovasculature, their functional relationship with arteriolar dynamics remains unclear. In this study, we established an *in vivo* two-photon imaging approach to simultaneously measure Ca^2+^ signals in cholinergic axonal varicosities and arteriolar diameters in urethane-anesthetized mice. An adeno-associated virus (AAV) vector (rAAV-ChAT-jGCaMP8s) was injected into the nucleus basalis of Meynert. *In vivo* imaging of the frontal cortex revealed bead-shaped GCaMP signals around the arterioles. Pinch stimulation transiently increased Ca^2+^ signals in periarteriolar varicosities, followed by arteriolar dilation, with an approximately 2-s delay between their peaks. Linear regression analysis disclosed a significant relationship between the magnitudes of these changes. This approach enabled simultaneous evaluation of cholinergic axonal activity and arteriolar dynamics *in vivo*, providing a tool to investigate the cholinergic regulation of cerebrovasculature.

**Highlights:**

- AAV-ChAT-GCaMP enables selective imaging of cholinergic projections
- Two-photon imaging reveals bead-shaped Ca^2+^ signals around arterioles
- Sensory stimulation increases periarteriolar cholinergic axonal Ca^2+^ signals
- Axonal Ca^2+^ signals are associated with arteriole dilation

## Introduction

Cholinergic neurons in the basal forebrain constitute a crucial neural system responsible for cognitive function. These neurons project axons to widespread brain regions such as the cerebral cortex, hippocampus, and olfactory bulb [1, 2]. These neurons release acetylcholine (ACh), which contributes to attention, learning, and memory as a neuromodulator [3]. Clinical studies revealed the loss of neurons in the nucleus basalis of Meynert (NBM) within the basal forebrain in patients with Alzheimer’s disease [4, 5]. Thus, cholinergic neurons remain important targets of investigation [6].

In addition to their neuromodulatory role, cholinergic neurons in the basal forebrain help regulate cerebral blood flow as arteriole dilators [7-11]. The cholinergic system is involved in cerebral blood flow regulation in response to somatosensory stimulation [12-15]. For example, skin pinch stimulation excites approximately 80% of basal forebrain neurons [16]. Additionally, pinch stimulation increases cerebral ACh release [17] and blood flow [18]. Such somatosensory stimulation-induced cerebral blood flow increases persist even after suppressing pressor responses, but they are attenuated by pharmacological blockade of ACh receptors [19-21]. Based on these findings, it has been proposed that skin stimulation excites cholinergic neurons in the basal forebrain, increases ACh release, and dilates the arterioles.

In cholinergic neurons, ACh is released from varicosities, which are bead-shaped axonal regions containing synapse vesicles [22-24]. The activity of cholinergic varicosity is monitored *in vivo* by two-photon imaging of Ca^2+^ signals using a genetically encoded Ca^2+^ sensor (i.e., GCaMP) expressed in basal forebrain cholinergic neurons via adeno-associated virus (AAV) vectors [25-28]. Additionally, a study using these methods revealed that an increase in axonal Ca^2+^ signaling precedes ACh release [29]. Histologically, the axonal varicosities of basal forebrain cholinergic neurons are distributed around blood vessels in the cerebral cortex [24]. However, the functional relationship between cholinergic axonal Ca^2+^ signals at periarteriolar varicosities and arteriolar responses has not been directly examined. Thus, we hypothesized that simultaneous *in vivo* imaging of arterioles and periarteriolar cholinergic axonal varicosity could permit examination of the contribution of basal forebrain cholinergic neurons to arteriolar response to somatosensory stimulation in a single experiment setting.

In the present study, we established an *in vivo* imaging method to simultaneously measure cerebral arteriolar diameter and Ca^2+^ signals in periarteriolar cholinergic axonal varicosities by expressing GCaMP in basal forebrain cholinergic neurons using an AAV vector carrying the choline acetyltrasnferase (ChAT) promoter [30] and labeling blood plasma using fluorescent dye [9, 31]. Using this approach, we further examined the relationship between periarteriolar varicosity activity and arteriolar dilation in response to skin pinch stimulation.

## Materials and Methods

### Animals

Seven male C57BL/6J mice (9–12 months old; weight, 28.3–30.9 g) were used. The animals were purchased from Japan Charlse River (Yokohama, Japan) and housed at the Tokyo Metropolitan Institute for Geriatrics and Gerontology. They were housed in individually ventilated cages with free access to a commercial pelleted diet (CRF-1, Oriental Yeast Co., Ltd., Tokyo, Japan) and filtered tap water containing 2 ppm chlorine. The vivarium was maintained at 22°C ± 1°C and 50% ± 10% relative humidity with a 12-h/12-h light/dark cycle (lights off at 20:00 h). The study protocol was approved by the Animal Care and Use Committee (approval number: 22020) and the committee for recombinant DNA experiments (approval number: 188) of the Tokyo Metropolitan Institute for Geriatrics and Gerontology. The protocol additionally followed the “Guidelines for Proper Implementation of Animal Experiments” established by the Japan Society for the Promotion of Science in 2006.

### Pre- and postoperative care for AAV vector injection

To ensure acclimatization to the breeding environment after surgery, gel food (MediGel^®^ Sucralose, ClearH_2_O, Westbrook, ME, USA), a heating pad (Kowa, Tokyo, Japan), and enrichment devices (wooden blocks and paper tunnel) were placed in cage for 3–4 days before surgery.

After the surgery, mice were housed individually until experimentation. The heating pad was kept in cage for the first 2 days after surgery, and gel food containing analgesic (meloxicam, 5 mg/kg/day) was provided alongside regular chow and drinking water for the first week. The enrichment devices were provided through the postoperative period, and the bedding was exchanged every week.

### Surgery for AAV vector injection

To express fluorescent protein in cholinergic neurons originating from the NBM, an AAV vector carrying ChAT promoter [30] was injected into the NBM. rAAV-ChAT-jGCaMP8s was injected for two-photon imaging in six mice (AAV9 serotype, titer = 6.05 × 10^12^ genome copies/mL, BC-4826, packaged by Brain Case, Wuhan, China).

Aseptic surgery was performed. Animals were anesthetized with isoflurane (initially 4% with room air, followed by 2%–3% during surgery). Prior to any surgical procedures, antibiotics (cefazoline sodium, 50 mg/kg) and analgesics (bupurenorphine, 0.03 mg/kg; meloxicam, 5 mg/kg) were subcutaneously injected. The scalp fur was trimmed using conventional clippers and disinfected using povidone–iodine solution (Isodine^®^, Shionogi Co., Ltd., Osaka, Japan) and 70% isopropyl ethanol. Ointment was applied to the eyes to prevent drying during surgery.

Each animal’s head was fixed to a stereotaxic instrument with ear bars (SR-5 M-S, Narishige, Tokyo, Japan) after local anesthetics were applied to the external ear canals (2% Xylocaine^®^ jelly, Sandoz Pharma K.K., Tokyo, Japan). The scalp was incised along the midline after procaine hydrochloride was subcutaneously injected. The skull was exposed and partly excised using a dental drill, and the dura was punctuated using a 30G syringe needle. The needle of a Neuros Syringe (7001 KH Neuros Syringe, HAMILTON, Reno, NV, USA) filled with the AAV vector was inserted into the cerebral cortex and slowly moved (10 μm/s) to the right NBM (anteroposterior, −0.9 mm from bregma; lateral, 2.2 mm from midline; ventral, 4.2 mm from the dorsal brain surface; [32]). The needle was further moved to 4.6 mm and slowly returned to the target position. The AAV vector was injected over 18 min at 50 nL/min (total volume, 900 nL) using an infusion pump (UMP3 UltraMicroPump and Micro4 Controller, World Precision Instruments, Sarasota, FL, USA). After the injection was completed, the needle was kept in place for 10 min and then slowly removed. The exposed cortical surface was sealed with a piece of plastic wrap (AsahiKASEI, Tokyo, Japan) and covered with dental cement (Inosit Baseliner, DMG Chemisch-Pharmazeutische Fabrik GmbH, Hamburg, Germany). The skin was sewn with a sterile 4-0 nylon suture and fixed with a tissue adhesive (Aron Alpha A, Daiichi Sankyo, Tokyo, Japan). Following surgery, warm saline (500 μL) was subcutaneously injected for body fluid supplementation.

In a preliminary experiment, we examined *via* immunohistochemistry whether an AAV vector carrying the ChAT-promoter could express a fluorescent protein in cholinergic axon varicosities in the cerebral cortex. A vector was injected into the right NBM in one mouse (AAV9 serotype, rAAV-ChAT-eGFP, titer = 5.43 × 10^12^ genome copies/mL, BC-4827, packaged by Brain Case). Three weeks after injection (Figure 1A), the animal was euthanized with isoflurane and transcardinally perfused with PBS and 4% paraformaldehyde in PBS using a peristatic pump. The brain was then removed. Cryosections (40 μm thick) were immunohistochemically stained (details are provided in the Supporting Information). Figure 1B presents confocal images of the frontal cortex. eGFP-expressing axons were sparsely observed, and the eGFP signals were colocalized with ChAT immunostaining, suggesting that the AAV vector carrying the ChAT promoter drives fluorescent protein expression in cholinergic axons.

**Figure 1.**
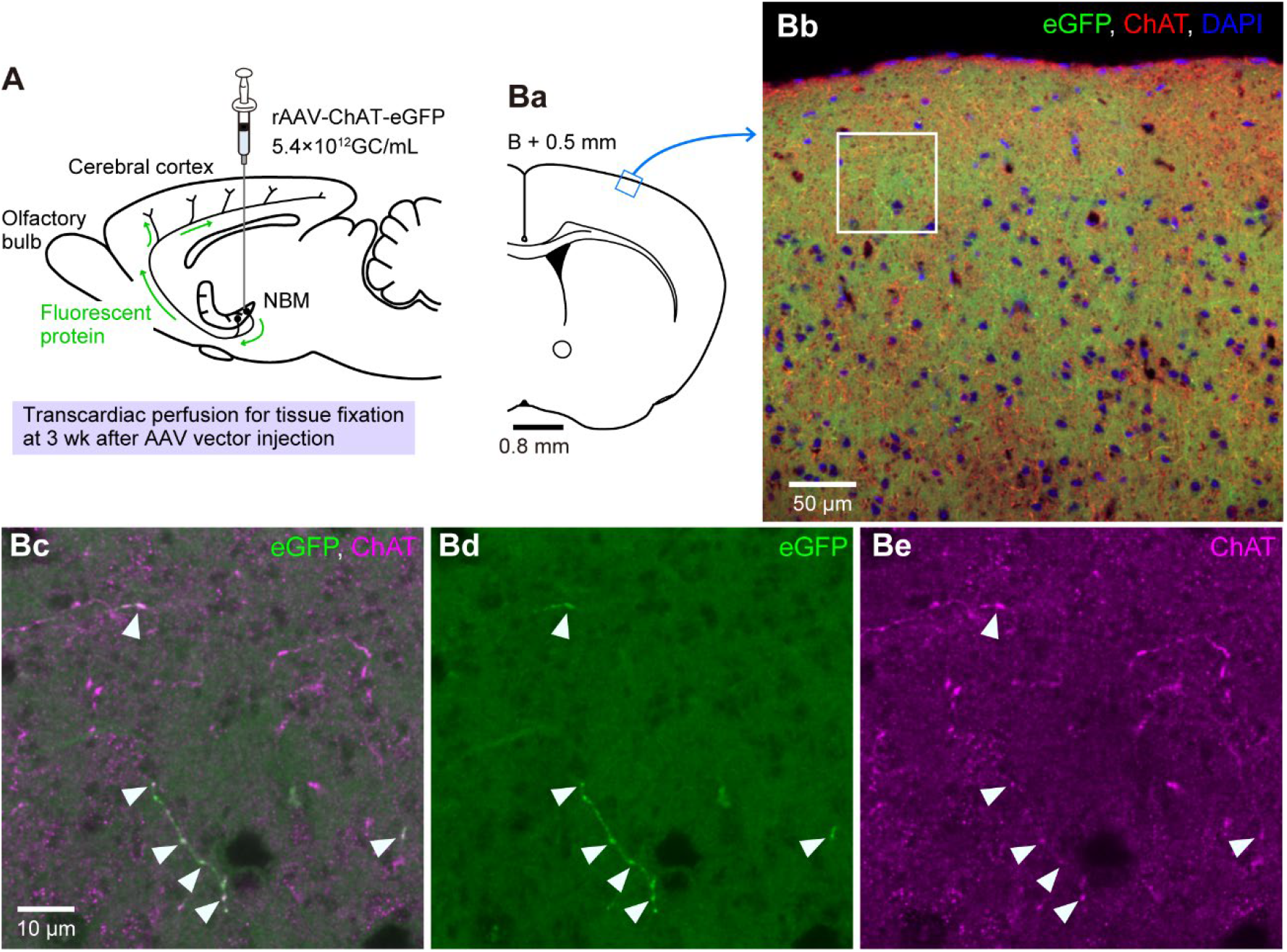
Colocalization of axonal eGFP with ChAT immunoreactivity in the mouse frontal cortex following rAAV-ChAT-eGFP injection into the NBM. **A**: Schematic diagram of AAV vector injection into the NBM. Animals were transcardially perfused 3 weeks after injection, allowing sufficient anterograde transport of the fluorescent protein into axons originating from the NBM. **B**: Confocal images showing eGFP-expressing axons originating from the NBM, ChAT immunoreactivity (anti-ChAT antibody), and DAPI staining in the frontal cortex. (a) Schematic diagram of a coronal section indicating the confocal imaging area. The diagram was drawn based on mouse brain atlas [32]. (b) Confocal image of the frontal cortex, as indicated by the light blue boxed region in (a). (c-e) Enlarged views of the white boxed region in (b). Arrowheads indicate colocalization of eGFP and ChAT immunoreactivity.

### In vivo two-photon imaging (GCaMP and arterioles)

*In vivo* two-photon imaging was performed 3–4 weeks after AAV vector injection (Figure 2A). Mice were first anesthetized with isoflurane (4% for approximately 3 min) and then anesthetized with urethane (1.4 g/kg, subcutaneously). Sufficient depth of anesthesia was determined based on loss of corneal and withdrawal reflexes, and additional anesthetic was administered as needed (injecting 5%–10% of the initial dose of urethane, or inhaling isoflurane). The trachea was cannulated, and the animals were artificially ventilated (MiniVent Type 845, Harvard Apparatus, MA, USA). Rectal temperature was maintained at 37.0°C–37.5°C using a feedback-controlled heating pad (BWT-100A, Bioresearch Center, Nagoya, Japan), and warm saline was continuously administered through a subcutaneously implanted 30G needle at 3 μL/min with an infusion pump (ESP-64, EICOM, Kyoto, Japan).

**Figure 2.**
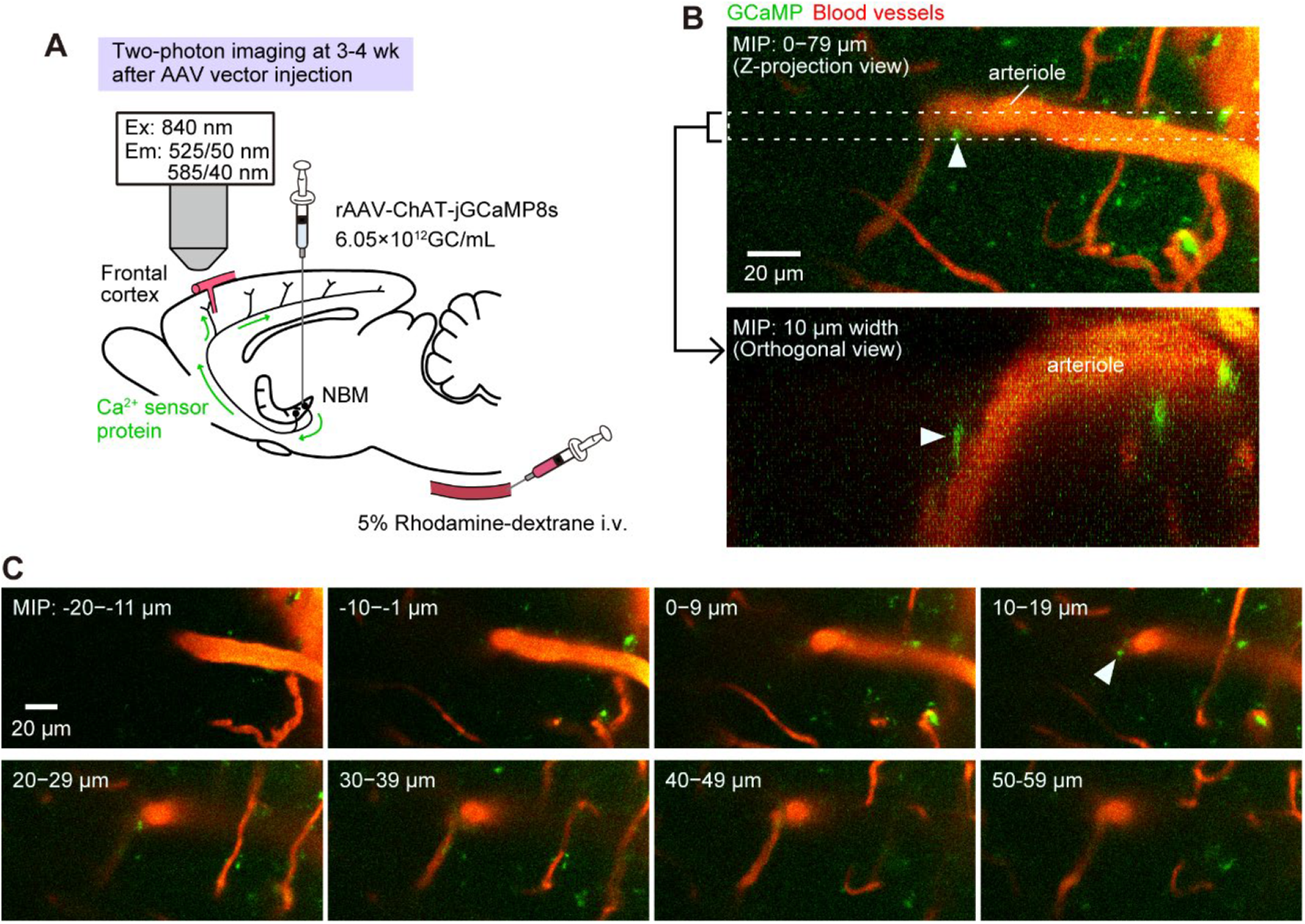
Three-dimensional *in vivo* two-photon imaging of cholinergic axonal Ca^2+^ signals and penetrating arterioles in the frontal cortex. **A**: Schematic diagram for *in vivo* two-photon imaging. Two-photon imaging of the frontal cortex was performed 3–4 weeks after AAV vector injection into the NBM, allowing sufficient anterograde transport of the Ca^2+^ sensor protein into axons of NBM cholinergic neurons. A fluorescent dye (5% rhodamine–dextran) was intravenously injected to visualize blood vessels. **B**: Example of three-dimensional two-photon imaging. The upper panel shows a Z-projection view with a maximum-intensity projection (MIP) constructed from z-stacks spanning 80 μm, acquired at 1-μm intervals. The lower panel shows an Orthogonal view, constructed from the region enclosed by the dashed line in the upper panel and displayed as a MIP over a 10-μm width. Arrowheads indicate a periarteriolar varicosity of a cholinergic axon. **C**: Series of z-stack images shown every 10 slices with a MIP.

Each animal’s head was fixed to a stereotaxic instrument with ear bars. A cranial window (approximately 2 mm in diameter) was made on the frontal cortex by thinning the skull using a dental drill, followed by soaking 10% disodium ethylenediamine tetraacetic acid solution for 10-15 min [31]. Dental cement (Fuji LUTE, GC Corp., Tokyo Japan) was applied around the cranial window to retain saline for immersing the objective lens. Rhodamine B isothiocyanate–dextran (molecular weight, 70 kDa; Sigma-Aldrich, St. Louis, MO, USA) dissolved in saline (5% solution) was intravenously administered before two-photon imaging at a volume of 80–100 μL. Fluorescence was excited using a two-photon laser (840 nm; Chameleon Vision II, Coherent, Santa Clara, CA, USA) with a ×25 correction collar lens (numerical aperture = 1.00, Leica Microsystems, Wetzlar, Germany). The emission signal was detected on an external detector through bandpass filters (525/50 and 585/40 nm) in the photon counting mode.

The site at which varicosities expressing detectable GCaMP signals were present near an arteriole branching from a pial arteriole originating from the anterior cerebral artery was selected for imaging. To obtain structural information, three-dimensional (*XYZ*) images were obtained. One *XY* plane image consisted of 1024 × 512 or 1024 × 1024 pixels with a pixel size of 0.2 μm, and depth scanning (*Z*) was performed with a step size of 1 μm.

To examine the influence of somatosensory stimulation (pinch stimulation of the left hindpaw), time-series images (*XYt*) were acquired for 30 s at a sampling rate of 3.8– 7.5 Hz. Pinch stimulation was applied for 10 s. The same investigator manually performed the stimulation and attempted to apply a consistent force (approximately 5–6 kg, measured using a Jamar® hydraulic pinch gauge, 5030HPG, J.A. Preston, Jackson, MI, USA). Trials were separated by at least 3 min.

### Analysis of in vivo two-photon imaging data

For *XYZ* imaging data, images were analyzed using Imaris software (v 10.0.0, Bitplane, Belfast, UK). The volume and minor radius of cholinergic varicosities and the distance from the closest arteriole were measured using the Surface function of the software. GCaMP signals and arterioles were smoothed using a Gaussian filter (diameters of 1 and 3 μm, respectively). The binarization threshold was determined by the full width at half-maximal intensity.

For *XYt* imaging data, the fluorescence intensity of GCaMP was measured using the Spots function in Imaris. Raw fluorescence (F) was extracted using a region of interest with a diameter of 3–4 μm that sufficiently included each varicosity. Relative fluorescence was calculated as ΔF/F(%) = [(F − F_0_)/F_0_] × 100. F_0_ was estimated by curve fitting to account for gradual baseline drift using exponential, polynomial, or power approximation functions, depending on the curvature before pinch stimulation. The diameter of the arteriole was measured using the Surface function in Imaris or ImageJ software with a plug-in program (Diameter, v 1.0). A 3-μm-diameter Gaussian filter was applied for smoothing. Imaris software was used when binarized arteriole images exhibited an oval shape, whereas ImageJ was used when the images had a non-oval shape. The binarization threshold was determined by the full width at half-maximal intensity.

The obtained values were temporally smoothed over three points and resampled at 3.8 Hz using Spike 2 software (v10.2.8., Cambridge Electronic Design, Cambridge, England).

### Statistical analysis

Statistical analyses were performed using Prism software (version 9.5.1, GraphPad Software Inc., Boston, MA, USA). The times to peak GCaMP signals and arteriolar diameter were compared using the paired t-test based on the normality of data, as tested using the Shapiro–Wilk test. The relationship between the magnitudes of the GCaMP signal and arteriolar diameter changes was evaluated using simple linear regression analysis. p < 0.05 indicated statistical significance. Data are expressed as the range (minimum-maximum) or median [interquartile range (IQR)].

## Results and Discussion

### Two-photon imaging of GCaMP signals in the varicosities of cholinergic axons and blood vessels in the frontal cortex

We explored the cerebral parenchyma and arterioles to observe GCaMP signals in proximity to arterioles in the frontal cortex using two-photon microscopy (Figure 2A). An example of three-dimensional imaging is presented as a maximum-intensity projection in Figure 2B and C. Bead-shaped GCaMP signals were found around the arterioles (arrowheads in Figure 2B and C), near capillaries, and within the cerebral parenchyma. Orthogonal views (lower panel of Figure 2B) and a series of horizontal images along the *Z*-axis (Figure 2C) indicated that a periarteriolar bead-shaped GCaMP signal was located approximately 15 μm below the cortical surface.

Among six mice, five completed the two-photon imaging experiment, whereas one mouse died during the experiment (cause unspecified). At least one periarteriolar bead-shaped GCaMP signal was detected in each of the five surviving mice. In total, seven bead-shaped GCaMP signals were detected (Table 1). Of these seven bead-shaped GCaMP signals, five were observed around penetrating arterioles at a depth of 5–40 μm from the cortical surface, and two were found around an arteriole at the cortical surface level before entering the brain parenchyma. These GCaMP signals were located at 0.33– 4.89 μm from the closest arteriole (lumen diameter, 9–16 μm).

**Table 1.** Profile of each periarteriolar varicosity of cholinergic axons.

| Varicosity # | Mouse # | Varicosity diameter ($\mu\text{m}$) | Depth from the cortical surface ($\mu\text{m}$) | Distance between varicosity and arteriole ($\mu\text{m}$) | Arteriolar diameter ($\mu\text{m}$) |
| --- | --- | --- | --- | --- | --- |
| 1 | M#1 | 1.97 | 40 | 2.79 | 13.8 |
| 2 | M#1 | 4.24 | 12 | 4.89 | 15.9 |
| 3 | M#2 | 2.77 | 13 | 0.33 | 12.9 |
| 4 | M#3 | 4.71 | 0 | 0.93 | 13.8 |
| 5 | M#3 | 4.17 | 0 | 4.21 | 11.3 |
| 6 | M#4 | 2.31 | 15 | 2.37 | 13.4 |
| 7 | M#5 | 3.3 | 5 | 2.25 | 9.1 |

Considering the thickness of the arteriolar wall (approximately 10% of the lumen diameter; [33]), the bead-shaped GCaMP signals were located within 3 μm of the outer surface of the arteriolar wall, consistent with perivascular varicosities reported in prior histological studies [24, 34]. Thus, the bead-shaped GCaMP signals observed in the present study represent periarteriolar varicosities of cholinergic axons originating from the NBM.

Fluorescence intensity changes in response to hindpaw pinch stimulation were analyzed in seven periarteriolar varicosities obtained from five mice. The binarization threshold was determined by the full width at half-maximal intensity for both cholinergic varicosities and arterioles. For the distance between varicosity and arteriole, the shortest distance between the surface of a cholinergic varicosity and the lumen of an arteriole was measured.

### Temporal relationship between cholinergic varicosity Ca^2+^ signals and arteriole diameter changes in response to pinch stimulation

We examined changes in GCaMP fluorescence intensity in the periarteriolar varicosities of cholinergic axons and arteriolar diameter in response to pinch stimulation (Figure 3A). Pinch stimulation was applied for 10 s following a 10-s pre-stimulation (baseline) period (Figure 3B, C). The GCaMP fluorescence transiently increased approximately 1 s after the onset of stimulation. The arteriole diameter increased and peaked approximately 3 s after the onset of stimulation before gradually declining toward the baseline level.

**Figure 3.**
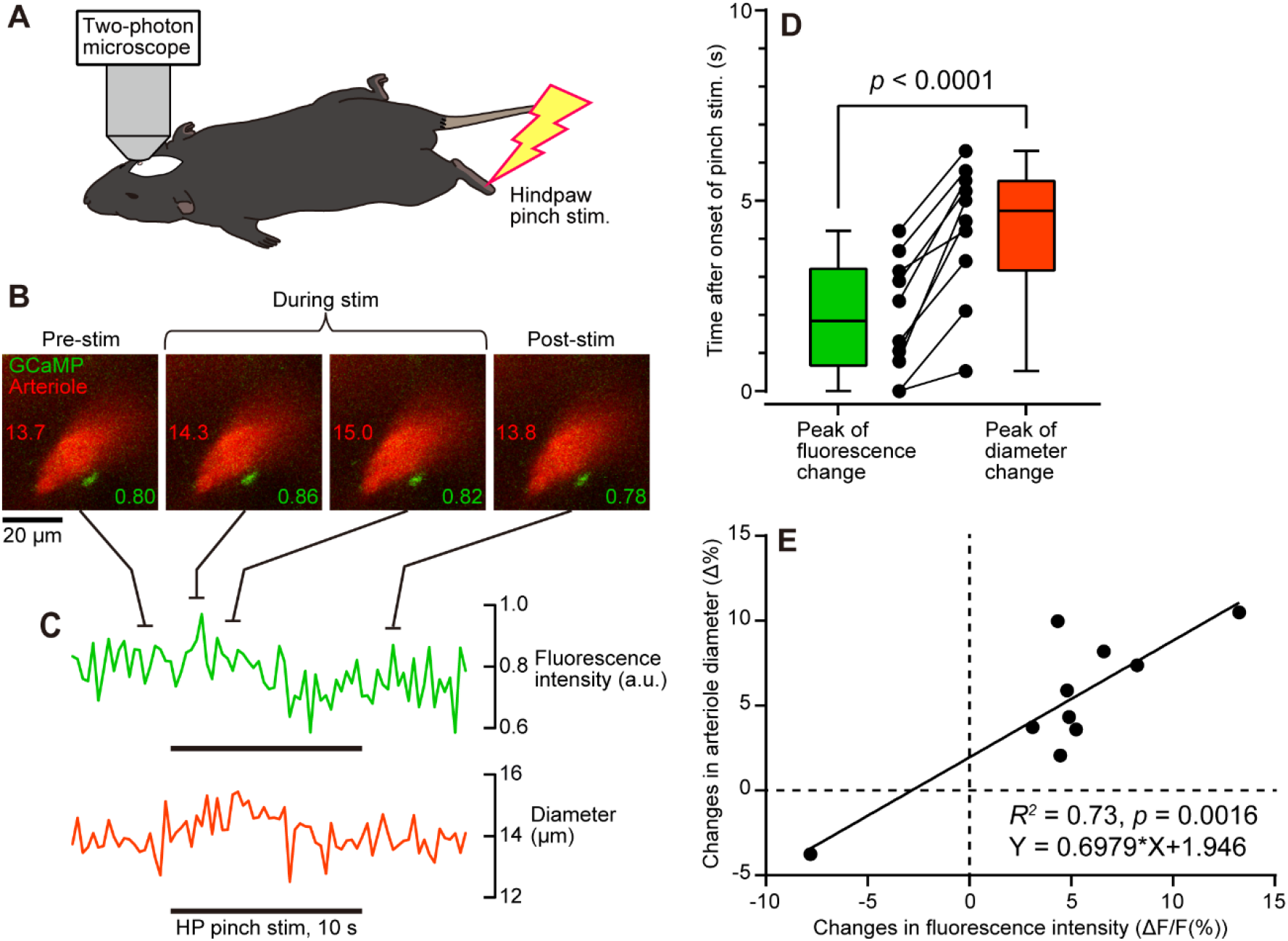
Relationship between cholinergic varicosity Ca^2+^ signals and arteriole diameter changes in responses to pinch stimulation. **A**: Schematic diagram of pinch stimulation applied to the hindpaw during *in vivo* two-photon imaging. **B**: Example images of a penetrating arteriole and periarteriolar GCaMP-expressing varicosity (average of three consecutive images each). Numerical values shown on each image are the fluorescent intensity (a.u., arbitrary units; in green) and luminal diameter (μm; in red). **C**: Time course of changes in GCaMP fluorescence intensity and arteriole luminal diameter measured (3.8 Hz of temporal resolution). Pinch stimulation was applied to the left hindpaw (HP) for 10 s. The thick horizontal line indicates the period of pinch stimulation. **D**: Temporal relationship between GCaMP signal and arteriolar diameter changes. The time to peak after the onset of pinch stimulation was compared using the paired t-test. Individual closed circles and lines indicate data obtained from each trial. A box represents the median and the 25%–75% range and whiskers indicate the minimum and maximum values. **E**: Relationship between the amplitudes of GCaMP fluorescent intensity and arteriole diameter changes (ΔF/F(%) and Δ%, respectively) was analyzed by linear regression.

Figure 3D summarizes the temporal relationship between GCaMP fluorescence and arteriolar diameter changes (n = 10 trials; 1–2 trials/varicosity). The peak GCaMP fluorescence occurred shortly after the onset of pinch stimulation [1.8 s (IQR, 0.6–3.3 s)], whereas the peak of arteriole diameter change occurred later at 4.7 s (IQR, 3.1–5.6 s). The paired t-test revealed that the peak of the GCaMP fluorescence change preceded the peak of the arteriole diameter change by 2.1 s (IQR, 1.8–3.2 s; p < 0.0001).

A similar temporal relationship between NBM activation and arteriolar dilation has been reported previously [9], supporting the validity of the present findings. This approach can simultaneously measure periarteriolar cholinergic varicosity activity and arteriole diameter changes in response to somatosensory stimulation, which had previously been studied separately [16-18], in a single experiment setting.

### Relationship between the amplitudes of cholinergic varicosity Ca^2+^ signals and arteriole diameter changes

Figure 3E summarizes the relationship between the amplitude of the GCaMP fluorescence intensity signal and arteriole diameter changes in response to pinch stimulation. GCaMP fluorescence increased following stimulation, excluding one trial in which GCaMP fluorescence decreased. Arteriole diameter changes were concordant with the GCaMP fluorescence intensity changes.

Simple linear regression analysis illustrated that changes in GCaMP fluorescence in response to pinch stimulation were significantly associated with arteriole diameter changes (R^2^ = 0.73, p = 0.0016; Figure 3E). This relationship between axonal activity and arteriolar dynamics might be supported by NBM stimulation frequency-dependent increases in cerebral ACh release and blood flow [10, 35, 36].

Changes in cholinergic varicosity activity around the penetrating arterioles following somatosensory stimulation are reasonable from the perspective of cerebral blood flow control. The penetrating arterioles represent a bottleneck in blood perfusion to the capillary network [37]. Therefore, periarteriolar cholinergic varicosities might contribute to cerebral blood flow regulation by controlling these arterioles.

## Conclusion

In conclusion, we established an experimental approach for simultaneously assessing the activity of periarteriolar cholinergic axonal varicosities and arteriolar dynamics *in vivo* by expressing GCaMP in basal forebrain cholinergic neurons using a ChAT promoter-driven AAV vector. Furthermore, using this approach, we examined the functional relationship of these structures in response to somatosensory stimulation. Although this approach revealed a temporal and quantitative association between periarteriolar cholinergic axonal varicosities and arteriolar diameter changes, further studies are required to determine causality.

Additionally, this experimental approach might represent a useful tool for investigating the mechanisms of cerebral blood flow regulation mediated by basal forebrain cholinergic neurons. Because we employed a ChAT promoter-driven AAV vector to express a Ca^2+^ sensor protein (jGCaMP8s) in cholinergic neurons of the basal forebrain in wild-type animals, this approach could be applicable beyond specific animal strains.

## Acknowledgements

The present study was partly supported by Tsumura & Co., Japan. The authors thank Prof. Fusao Kato and Dr. Yukari Takahashi at The Jikei University School of Medicine for valuable advice on the use of AAV vectors, and Dr. Shuichi Yanai and Ms. Tomoko Arasaki at the TMIG Animal Facility for their technical support.

