## Supplemental information of methods for "Simultaneous *in vivo* imaging of Ca^2+^ signals in periarteriolar cholinergic axonal varicosities and arteriole diameter changes in the mouse cerebral cortex"

**Supporting Information**

**Methods**

***Histological analysis***

The removed brain was postfixed in 4% paraformaldehyde in PBS overnight at 4°C and cryoprotected in sucrose solution (10% sucrose in PBS for 4 h and 20% sucrose in PBS for 24–48 h). After rinsing in PBS, the brain was embedded in OCT compound in a plastic mold (Tissue-Tek^®^, Sakura Finetek Japan Co., Ltd., Tokyo, Japan; Square-S22, Polysciences, Inc., Warrington, PA, USA) and frozen on an aluminum block with liquid nitrogen. The tissue was stored at −80°C until further processing.

The brain was sectioned coronally at a thickness of 40 μm using a cryostat (CM3050S, Leica Microsystems, Wetzlar, Germany) and placed on glass slides. After drying, sections were washed with PBS and blocked in PBS containing 0.1% Triton X-100 and 3% normal donkey serum for 1 h at room temperature. Sections were then incubated with a primary antibody (anti-choline acetyltransferase (ChAT), 1:100, AB144P, Millipore, Burlington, MA, USA; RRID: AB_90661) in blocking solution for 24 h at 4°C. After washing with PBS, the sections were incubated with a secondary antibody (Alexa Fluor 594, 1:200, Thermo Fischer Scientific, Waltham, MA, USA; RRID: AB_2534105) for 2 h at room temperature. After washing with PBS containing 0.02% NaN_3_, sections were mounted using a mounting medium containing 4′,6-diamidino-2-phenylindole (DAPI) (Vectashield plus H-2000, Vector Laboratories, Inc., Newark, CA, USA).

The immunohistochemically stained sections were imaged using a confocal microscope (STELLARIS WLL, Leica Microsystems) with a ×40 HC PL APO objective (NA =1.30, oil CS2), a digital zoom of ×0.75–3.88 and sequential line scanning for each channel. Colocalization of eGFP signals with anti-ChAT immunostaining was assessed.
